# Pharmaco-genetic screen to uncover actin regulators targeted by prostaglandins during *Drosophila* oogenesis

**DOI:** 10.1101/722355

**Authors:** Andrew J. Spracklen, Maureen C. Lamb, Christopher M. Groen, Tina L. Tootle

**Author notes:** Department of Biology, University of Massachusetts, Amherst, MA, 01003. Regenerative Neurobiology Laboratory, Mayo Clinic, Rochester, MN 55905.

## Abstract

Prostaglandins (PGs) are lipid signaling molecules with numerous physiologic functions, including pain/inflammation, fertility, and cancer. PGs are produced downstream of cyclooxygenase (COX) enzymes, the targets of non-steroidal anti-inflammatory drugs (NSAIDs). In numerous systems, PGs regulate actin cytoskeletal remodeling, however, their mechanisms of action remain largely unknown. To address this deficiency, we undertook a pharmaco-genetic interaction screen during late-stage *Drosophila* oogenesis. *Drosophila* oogenesis is as an established model for studying both actin dynamics and PGs. Indeed, during Stage 10B, cage-like arrays of actin bundles surround each nurse cell nucleus, and during Stage 11, the cortical actin contracts, squeezing the cytoplasmic contents into the oocyte. Both of these cytoskeletal properties are required for follicle development and fertility, and are regulated by PGs. Here we describe a pharmaco-genetic interaction screen that takes advantage of the facts that Stage 10B follicles will mature in culture and COX inhibitors, such as aspirin, block this *in vitro* follicle maturation. In the screen, aspirin was used at a concentration that blocks 50% of the wild-type follicles from maturing in culture. By combining this aspirin treatment with heterozygosity for mutations in actin regulators, we quantitatively identified enhancers and suppressors of COX inhibition. Here we present the screen results and initial follow-up studies on three strong enhancers – Enabled, Capping protein, and non-muscle Myosin II Regulatory Light Chain. Overall, these studies provide new insight into how PGs regulate both actin bundle formation and cellular contraction, properties that are not only essential for development, but are misregulated in diseases.

## Introduction

Many physiological functions, including pain/inflammation, reproduction, heart health/disease, and cancer, are mediated by lipid signals termed prostaglandins (PGs) (Tootle, 2013). PGs are produced at their site of action by cyclooxygenase (COX) enzymes, which are inhibited by non-steroidal anti-inflammatory drugs (NSAIDs). One cellular target of PGs is the actin cytoskeleton (Halbrugge *et al.*, 1990; Nolte *et al.*, 1991; Kawada *et al.*, 1992; Peppelenbosch *et al.*, 1993; Aszodi *et al.*, 1999; Pierce *et al.*, 1999; Banan *et al.*, 2000; Bearer *et al.*, 2002; Dormond *et al.*, 2002; Martineau *et al.*, 2004; Bulin *et al.*, 2005; Birukova *et al.*, 2007). However, the mechanisms by which PGs regulate actin remodeling remain largely unknown. To address this, we undertook a screen to identify the specific targets of PGs during *Drosophila* oogenesis.

*Drosophila* oogenesis provides a powerful model for uncovering the mechanisms by which PGs regulate actin remodeling (Spracklen and Tootle, 2015). Actin-dependent morphogenic events necessary for follicle development (Figure 1A) are regulated by the coordinated activity of numerous actin binding proteins [reviewed in (Hudson and Cooley, 2002)]. There are 14 stages of follicle development. Each follicle consists of 16 germline-derived cells – an oocyte and 15 support cells, termed nurse cells – and ∼1,000 somatically-derived follicle cells. During Stage 10B (S10B), cage-like arrays of parallel actin filament bundles rapidly extend from the nurse cell membranes toward the nuclei (Guild *et al.*, 1997; Huelsmann *et al.*, 2013). These bundles hold the nuclei in place during Stage 11 (S11), when the nurse cells undergo a rapid actomyosin-based contraction to transfer their cytoplasmic contents into the expanding oocyte in a process termed nurse cell dumping (Wheatley *et al.*, 1995). Nurse cell dumping requires PGs, as loss of the *Drosophila* COX-like enzyme Pxt blocks this process (Tootle and Spradling, 2008). Specifically, PGs are required for bundle formation, cortical actin integrity, and cellular contraction (Tootle and Spradling, 2008; Groen *et al.*, 2012).

**Figure 1.**
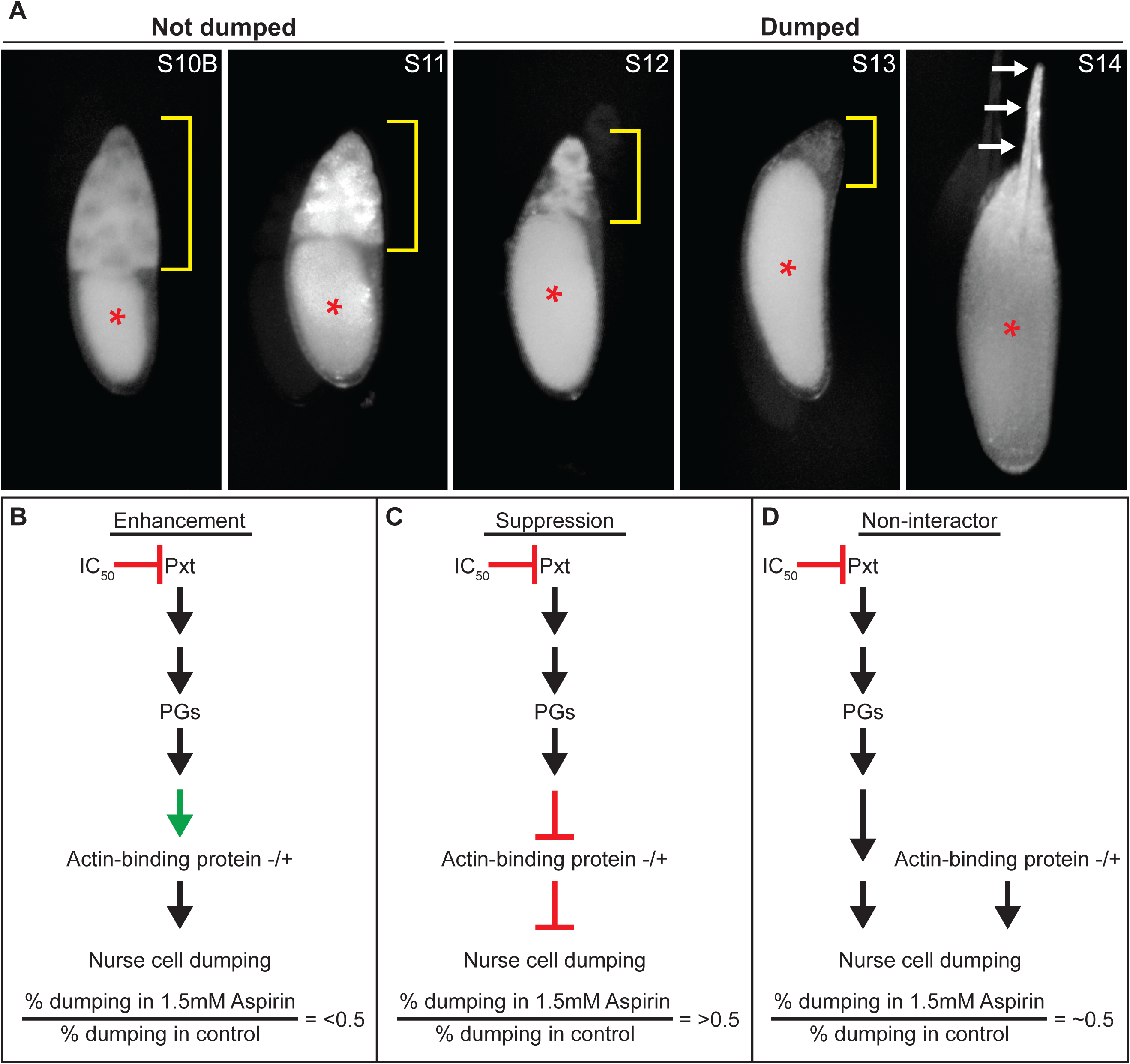
Examples of late-stage *Drosophila* follicles and screen rationale. A. Representative images of S10B-S14 *Drosophila* follicles taken using a stereo dissecting scope; anterior is at the top. Red asterisks indicate the oocyte, yellow brackets mark the nurse cells, and white arrows indicate the dorsal appendages. In the IVEM assay, follicles that remain in S10B-11 are considered undumped, whereas follicles that have reached S12-14 are considered dumped. B-D. Diagrams illustrating the rationale behind the pharmaco-genetic interaction screen. If a particular actin-binding protein is positively regulated by PG signaling to promote actin remodeling during S10B, then heterozygosity for a strong allele of that actin-binding protein would be expected to enhance follicle sensitivity to COX inhibition, resulting in an expected dumping index (% dumping in 1.5mM aspirin/%dumping in control) of less than 0.5 (B). If a particular actin-binding protein is negatively regulated by PG signaling to promote actin remodeling during S10B, then heterozygosity for a strong allele of that actin-binding protein would be expected to suppress follicle sensitivity to COX inhibition, resulting in an expected dumping index greater than 0.5 (C). Reduced levels of an actin-binding protein that is not a downstream target of PG signaling during S10B would not be expected to modify follicle sensitivity to COX inhibition, resulting in an expected dumping index of ∼0.5 (D).

Multiple lines of evidence indicate PGs directly regulate actin remodeling. First, mRNA levels of actin regulators are unchanged in *pxt* mutants (Tootle *et al.*, 2011). This has been verified at the protein level for a number of actin binding proteins (Groen *et al.*, 2012; Spracklen *et al.*, 2014). Second, pharmacologic disruption of PG signaling acutely disrupts actin remodeling (Tootle and Spradling, 2008), suggesting that PGs likely post-translationally regulate actin binding proteins to rapidly modulate actin remodeling. Together these data led us to hypothesize that PG signaling coordinates the activities of multiple actin regulators to promote both actin remodeling during S10B and cellular contraction during S11.

Here we present the results of a pharmaco-genetic interaction screen to identify actin binding proteins functioning downstream of PG signaling during S10B-11 of *Drosophila* oogenesis. This screen identified a number of actin regulators and interacting proteins including Capping protein (Cp; *Drosophila* Cpa and Cpb), E-Cadherin (*Drosophila* Shotgun, Shg), Enabled (Ena), Fascin (*Drosophila* Singed, Sn), Lamellipodin (*Drosophila* Pico), and the non-muscle Myosin II Regulatory Light Chain (MRLC; *Drosophila* Spaghetti squash, Sqh) as candidate downstream targets of PG signaling. Here, we present a summary of the screen results and follow-up studies on Ena, Cp, and MRLC.

## Methods

### Fly husbandry

Fly stocks were maintained at 21°C on standard cornmeal-agar-yeast food. Flies were fed with wet yeast paste daily and aged for 3-5 days for *in vitro* follicle maturation (IVEM) assays and ovary analyses, including immunofluorescence and immunoblotting. *y*^*1*^*w*^*1*^ (*yw*) was used as the wild-type control in experiments. The sources of stocks used in the pharmaco-genetic interaction screen are indicated in Table 1.

**Table 1:**
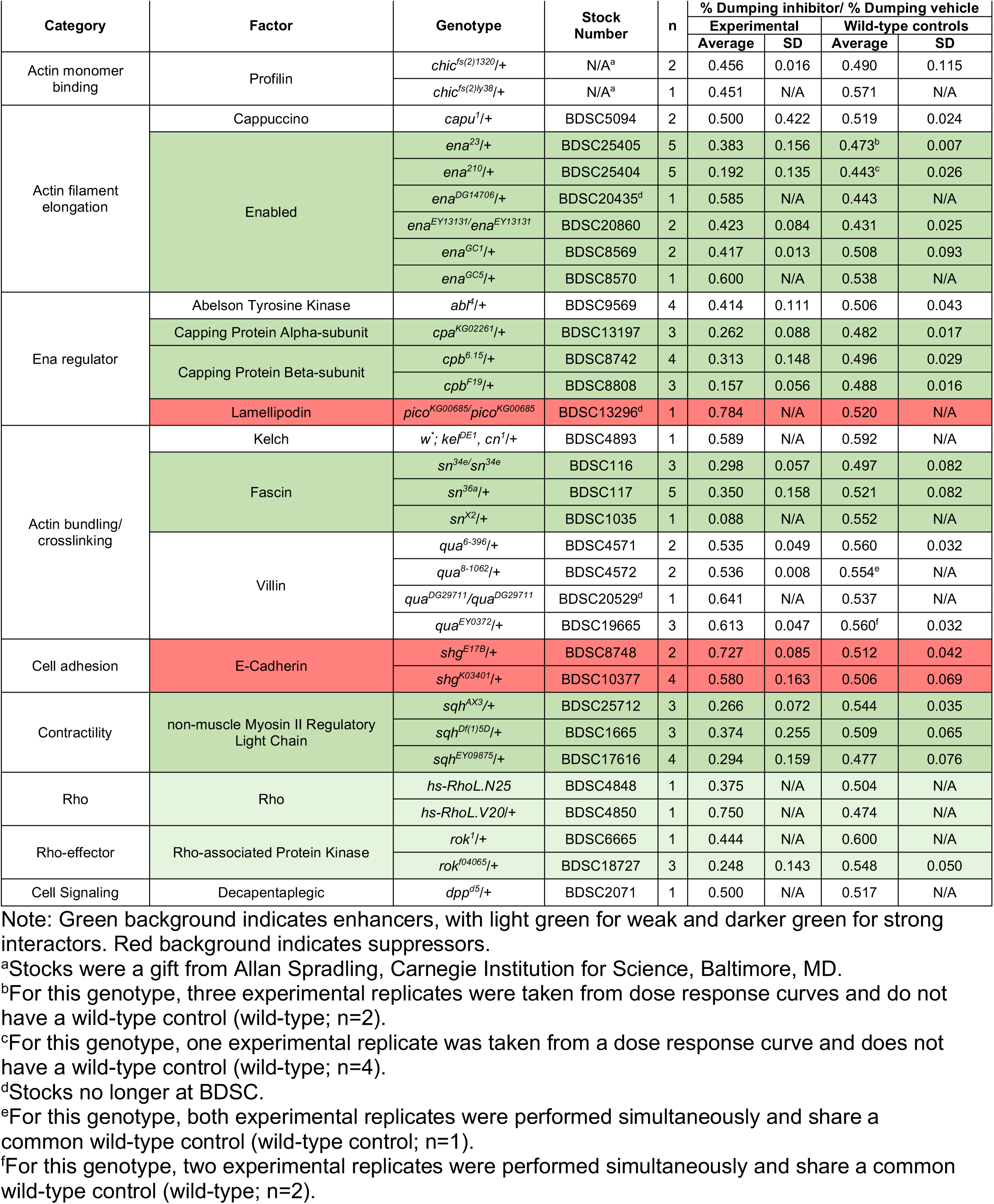
Average dumping indices for indicated genotypes tested in the pharmaco-genetic interaction screen and their respective wild-type controls.

### *In vitro* follicle maturation (IVEM)

IVEM assays were performed as previously described (Tootle and Spradling, 2008; Spracklen and Tootle, 2013). Briefly, for each genotype analyzed, whole ovaries were dissected out of mated adult females in room temperature IVEM media (10% heat inactivated fetal bovine serum (FBS) (Atlanta Biologicals, Flowery Branch, GA, USA), 1X penicillin/streptomycin (100X penicillin/streptomycin, Gibco, Life Technologies, Carlsbad, CA, USA), in Grace’s insect media (Lonza, Walkersville, MD, USA)) into a 9-well glass plate using forceps. Ovaries were immediately transferred to a clean well with fresh IVEM media and Stage 10B (S10B) follicles were isolated using dissecting needles mounted in pin vises and transferred to a clean well. Once 20-30 S10B follicles per condition were isolated, they were transferred to a clean well in a 24-well plate. Any IVEM media transferred with the follicles was gently aspirated using a pulled Pasteur pipette and immediately replaced with 1 mL of IVEM media containing vehicle only (3μl of 100% ethanol per 1 mL of IVEM media) or 1.5mM aspirin (3μl of 0.5M aspirin (Cayman Chemical, Ann Arbor, MI, USA) in 100% ethanol per 1 mL of IVEM media). Treated follicles were allowed to mature overnight at room temperature. The assay was scored by determining the percentage of follicles that completed nurse cell dumping in each condition. This was determined by assessing whether the follicles stalled at S10B or whether they progressed to Stage 11 (S11), 12 (S12), or 13/14 (S13/14) based on overall follicle morphology (Figure 1A). Follicles scored as S10B or S11 were considered to have not completed nurse cell dumping, while follicles that reached S12 or S13/14 were considered to have completed nurse cell dumping.

### Screen analysis

In order to compare genotypes across the screen, we calculated a dumping index (% follicles dumping in 1.5mM aspirin/% follicle dumping in control media) for each replicate. Average dumping indices and standard deviations for each experimental genotype and their corresponding controls are reported in Table 1. To aid in the identification of potential interactions in this screen, we focused on differences in the dumping index between experimental genotypes and wild-type controls by normalizing the data. To do this, we set the dumping index for wild-type controls to zero and normalized the corresponding experimental dumping indices by subtracting the wild-type dumping index from that of the experimental group for each individual experiment. The average normalized dumping index, and their standard deviation, are plotted in Figure 2. For the purpose of this screen, we arbitrarily defined strong interactors as those showing a response of greater than three standard deviations from wild-type controls, weak interactors as those showing a response between one and three standard deviations from wild-type controls, and non-interactors showing a response of less than one standard deviation from wild-type controls in their ability to modify sensitivity to COX inhibition. For statistical analyses, we used unpaired t tests to compare means between experimental and control groups. Graphs generated and statistical comparisons made using Prism8 (GraphPad Software, La Jolla, CA, USA).

**Figure 2.**
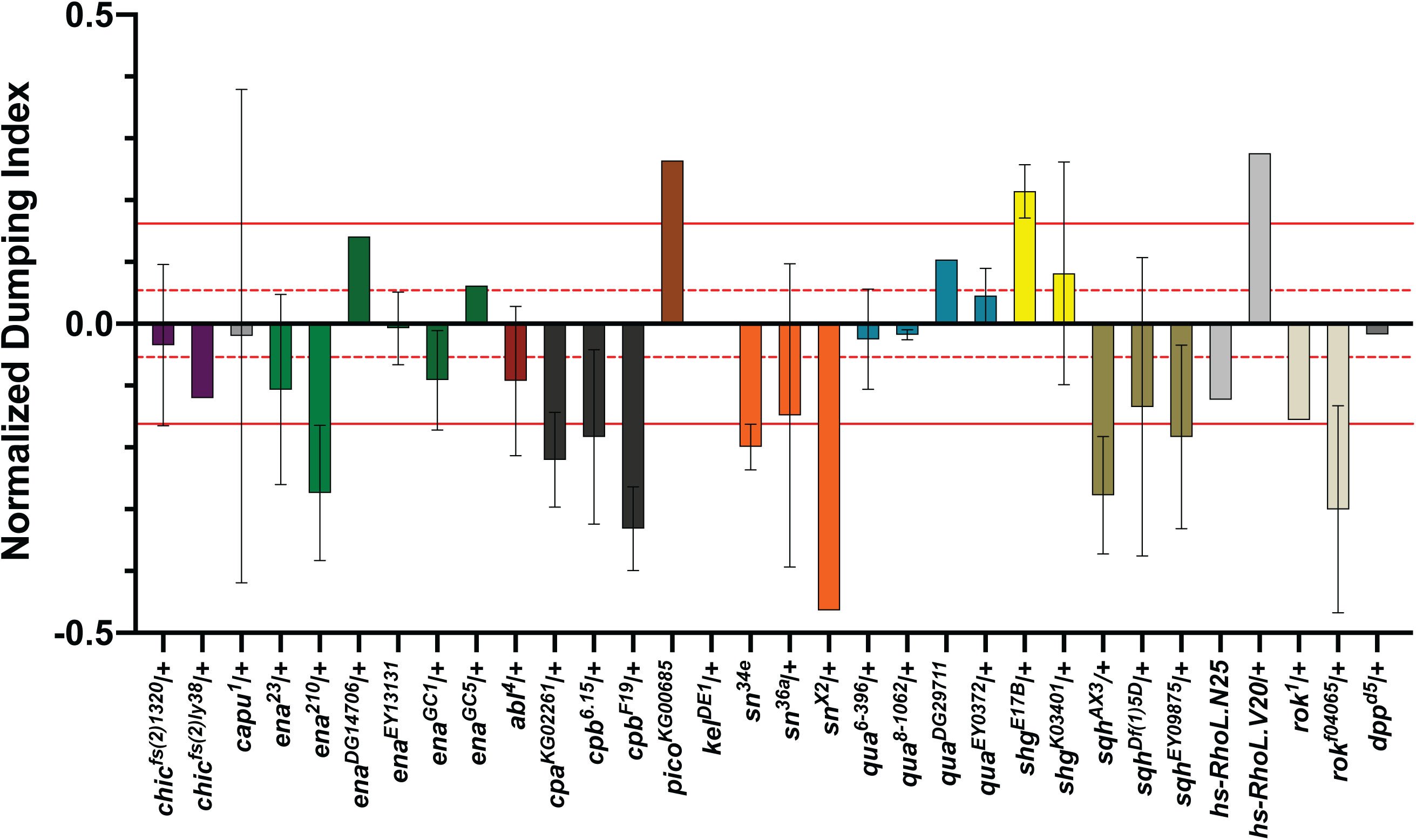
Pharmaco-genetic interaction screen reveals multiple actin- binding proteins as candidate downstream targets of prostaglandin signaling. Chart of the normalized dumping indices for all genotypes tested in the first pass of the pharmaco-genetic interaction screen. Wild-type control values were set to zero and normalized experimental dumping indices were calculated by subtracting the wild-type dumping index from the dumping index of the experimental group for each individual experiment. Dashed red lines indicate ±1 standard deviation (±0.054) from all wild-type control values. Solid red lines represent ±3 standard deviations (±0.162) from all wild-type control values. Genotypes exhibiting normalized dumping indices between 0 and ±1 standard deviation from wild-type control values are classified as non-interactors. Genotypes exhibiting normalized dumping indices falling between ±1 to ±3 standard deviations from wild-type control values are classified as weak interactors. Genotypes exhibiting normalized dumping indices falling outside of ±3 standard deviations from wild-type control values are classified as strong interactors. Error bars = standard deviation (SD).

### Western blots

Approximately 5 whole ovary pairs were dissected in room temperature Grace’s insect media (Lonza, Walkersville, MD, USA or Thermo Fisher Scientific, Waltham, MA), transferred to a 1.5ml microcentrifuge tube containing 50µL of Grace’s media, an equal volume of 2X SDS Sample Buffer was added and the tissue lysed by grinding with a plastic pestle. Western blots were performed using standard methods. The following primary antibodies were used: mouse α*−*Enabled (5G2) (Goodman, C.; obtained from the Developmental Studies Hybridoma Bank (DSHB), 1:200; α-Zipper (Karess, R.; Institut Jacques Monod, Paris, France; (Wheatley *et al.*, 1995)), 1:1000 and mouse α−αTubulin T9026 (Sigma-Aldrich, St. Louis, MO, USA), 1:5000. All blots had 0.1% Tween 20 added to the primary antibody in 5% milk diluted in 1× Tris-buffered saline. The following secondary antibodies were used: Peroxidase-AffiniPure Goat Anti-Mouse IgG (H+L), 1:5000 and Peroxidase-AffiniPure Goat Anti-Rabbit IgG (H+L), 1:5000 (Jackson ImmunoResearch Laboratories, West Grove, PA, USA). Blots were developed with SuperSignal West Pico or Femto Chemiluminescent Substrate (Thermo Scientific, Waltham, MA, USA) and imaged using the ChemiDoc-It Imaging System and VisionWorksLS software (UVP, Upland, CA, USA) or Amersham Imager 600 (GE Healthcare Life Sciences, Chicago, IL). Bands were quantified using densitometry analysis in ImageJ (Abramoff *et al.*, 2004). Ena levels were assessed using 2 independent biological samples. Zipper levels were assessed using a minimum of 3 independent, biological samples, and statistical significance was determined using a two-sample t-test with unequal variance in Excel (Microsoft, Redmond, WA, USA).

### Immunofluorescence

Whole ovaries were dissected in room temperature Grace’s insect media (Thermo Fisher Scientific, Waltham, MA). Active-MRLC staining was performed using a protocol provided by Jocelyn McDonald (Majumder *et al.*, 2012; Aranjuez *et al.*, 2016). Briefly, ovaries were fixed for 20 min at room temperature in 4% paraformaldehyde in 1X phosphate-buffered saline (PBS) and 0.2% Triton X-100. Samples were blocked by incubating in Triton antibody wash (1XPBS, 0.2% Triton X-100, and 5% bovine serum albumin [BSA]) for 30 min. Primary antibodies were incubated for at least 30 hours at 4°C. The rabbit anti-pMRLC (S19; Cell Signaling, Davers, MA) was diluted 1:125 in Triton antibody wash. Alexa Fluor 647–phalloidin (Invitrogen, Life Technologies, Grand Island, NY) was included with both primary and secondary antibodies at a concentration of 1:250. After six washes in Triton antibody wash (10 min each), the secondary antibody Alexa Fluor 488::goat anti-rabbit (Invitrogen, Life Technologies) was diluted 1:500 in Triton antibody wash and incubated overnight at 4°C. After six washes in Triton antibody wash (10 min each), 4’,6-diamidino-2-phenylindole (DAPI, 5 mg/mL) staining was performed at a concentration of 1:5000 in 1XPBS for 10 min at room temperature. Ovaries were mounted in 1 mg/ml phenylenediamine in 50% glycerol, pH 9 (Platt and Michael, 1983). All experiments were performed a minimum of three independent times.

### Image acquisition and processing

Confocal z-stacks of *Drosophila* follicles were obtained using Zen software on a Zeiss 700 LSM mounted on an Axio Observer.Z1 using a Plan-Apochromat 20x/0.8 working distance (WD) = 0.55 M27 (Carl Zeiss Microscopy, Thornwood, NY). Maximum projections (2-3 confocal slices), merged images, rotation, cropping, and pseudocoloring, including rainbow RGB intensity coloring, were performed using ImageJ software (Abramoff *et al.*, 2004).

### Data and reagent availability

Fly stocks and detailed protocols are available upon request. The authors affirm that all data necessary for confirming the conclusions of the article are present within the article, figures, and table.

## Results and Discussion

### Pharmaco-genetic interaction screen

To identify actin regulators and interacting proteins acting downstream of PG signaling during S10B of *Drosophila* oogenesis, a pharmaco-genetic interaction screen was undertaken. This screen took advantage of the following: 1) S10B follicles mature to S14 in *in vitro* culture conditions (Figure 1A; (Tootle and Spradling, 2008; Spracklen and Tootle, 2013)), 2) COX inhibitor treatment suppresses follicle maturation (Tootle and Spradling, 2008; Spracklen and Tootle, 2013), and 3) multiple mutant alleles of most actin regulators and interacting proteins are readily available. In this screen, we hypothesized that if a particular actin regulator were a downstream target of PG signaling, then reduced levels of that actin regulator (i.e., through heterozygosity for strong alleles or homozygosity for weak alleles) would enhance or suppress the inhibition of nurse cell dumping and subsequent follicle maturation due to COX inhibitor treatment. For example, if PG signaling positively regulates Protein X to promote actin remodeling during S10B, then heterozygosity for a strong allele of *protein x* would be expected to enhance follicle sensitivity to COX inhibition (Figure 1B). However, if PG signaling negatively regulates Protein Y to promote actin remodeling during S10B, then heterozygosity for a strong allele of *protein y* would be expected to suppress follicle sensitivity to COX inhibition (Figure 1C). In contrast, reduced levels of an actin binding protein that is not a downstream target of PG signaling during S10B would not be expected to modify follicle sensitivity to COX inhibition (Figure 1D).

Based on this rationale, we conducted the first pass of a pharmaco-genetic interaction screen using a concentration of aspirin (1.5mM) empirically determined to reproducibly inhibit ∼50% of a population of wild-type S10B follicles from completing nurse cell dumping in culture (Tootle and Spradling, 2008). Using this dose of aspirin ensured we could readily observe both suppression and enhancement of follicle sensitivity to COX inhibition. For each genotype tested, isolated S10B follicles were cultured overnight in either vehicle-treated or aspirin-treated media. We then determined the ratio of the percentage of follicles completing nurse cell dumping in aspirin treated media to the percentage of follicles completing nurse cell dumping in control media for each genotype analyzed (subsequently referred to as the dumping index). Follicles in S10B-11 were considered to have not completed dumping, while those in S12-14 were scored as completing nurse cell dumping (Figure 1A). In order to control for potential fluctuations in aspirin concentration (i.e., solvent evaporation) and any inter-experiment variability, wild-type (*yw*) controls were included in each experiment. Using this approach, we would expect wild-type follicles to exhibit a dumping index of ∼0.5, indicating half of the population of S10B follicles failed to complete nurse cell dumping in aspirin treated media. Experiments in which the wild-type dumping index was less than 0.4 or greater than 0.6 were excluded from further analyses. We observed an average dumping index of 0.505 (±0.054) for the inclusive data set of all wild-type controls (n=59). Aggregate data for all genotypes analyzed and their respective wild-type controls, including the number of trials (n), the average dumping index, and their standard deviation (SD), are summarized in Table 1.

### Identifying interactors

To aid in the identification of potential interactors in this screen, we wanted to focus on the differences in dumping index between wild-type control and experimental groups. To achieve this, the dumping index for wild-type controls were set to zero and normalized experimental dumping indices were calculated by subtracting the wild-type dumping index from the dumping index of the experimental group for each individual experiment. Normalized dumping indices for each genotype were then averaged and plotted (Figure 2). By normalizing the data in this manner, the plotted values represent the net change in dumping index due to reduced levels of the factor tested. Positive values indicate that reduction of the factor tested suppressed sensitivity to COX inhibition, while negative values indicate enhanced sensitivity. The magnitude of this change indicates the relative strength of the observed suppression or enhancement. Using this strategy, we classified a number of genes falling into one of three categories: non-interactors, weak interactors, and strong interactors.

Based on this normalized data set, we defined non-interactors as any genotype that fails to exhibit a change in dumping index greater than one standard deviation away from wild-type values (±0.054; calculated from non-normalized aggregate wild-type controls). Factors falling into this category are comprised of both actin binding proteins and signaling pathway components known to play critical roles throughout *Drosophila* oogenesis, including: the formin Cappuccino; the actin cross-linking protein Kelch; the actin monomer binding protein Profilin (*Drosophila* Chickadee, Chic); the actin bundling protein Villin (*Drosophila* Quail); a TGFβ-like and BMP-like ligand Decapentaplegic (Dpp); and Abelson Tyrosine Kinase (Abl) (Table 1, no background color, and Figure 2).

Genotypes exhibiting a change in dumping index between one and three standard deviations away from wild-type values were defined as weak interactors (between ±0.054 and ±0.162). Factors falling into this category include the small GTPase, Rho (*Drosophila* Rho-like, RhoL) and the Rho effector, Rho-associated Protein Kinase (ROCK, *Drosophila* ROK) (Table 1, light green background, and Figure 2).

Strong interactors were defined as any genotype exhibiting a change in dumping index equal to or greater than three standard deviations away from wild-type values (±0.162). Based on this definition, we identified a number of factors whose genetic reduction strongly sensitizes follicles to the effects of COX inhibition, including: the actin filament barbed end capper Cp; the actin elongation factor Ena; the contractile protein MRLC; and the actin bundling protein Fascin (Table 1, darker green background and Figure 2). Additionally, we uncovered two factors whose genetic reduction strongly suppresses follicle sensitivity to the effects of COX inhibition: the cell adhesion molecule E-Cadherin and the Enabled/Vasodilator-stimulated phosphoprotein (Ena/VASP) ligand Lamellipodin (Table 1, red background and Figure 2). Notably the strong and weak interactors represent actin binding proteins and related factors that participate in parallel actin bundle formation and nurse cell contractility – processes that fail when PG signaling is lost (Tootle and Spradling, 2008; Groen *et al.*, 2012; Spracklen *et al.*, 2014).

While all of the candidates uncovered in this screen are interesting, we chose to follow up on a subset of strong interactors. One such interactor is Fascin. Fascin is an actin bundling protein (reviewed in (Edwards and Bryan, 1995) required for parallel actin filament bundle formation during S10B (Cant *et al.*, 1994). Reduction of Fascin enhances sensitivity to COX inhibition (Table 1 and Figure 2). Through subsequent pharmacologic and genetic studies, we validated Fascin as a novel downstream target of PG signaling (Groen *et al.*, 2012). However, the mechanism(s) by which Fascin is regulated downstream of PG signaling remain poorly understood. Recent studies indicate PGs regulate the subcellular localization of Fascin (Groen *et al.*, 2015; Jayo *et al.*, 2016; Kelpsch *et al.*, 2016). Additionally, it is likely that PGs control Fascin through post-translational modifications (Groen and Tootle, unpublished data). Here we present further data for three strong interactors: Ena, Cp, and MRLC.

### Heterozygosity for a localization defective Ena allele sensitizes follicles to COX inhibition

Ena, the sole *Drosophila* homolog of the Ena/VASP family of actin elongation factors (Gertler *et al.*, 1995), is a large protein containing multiple domains including: an Ena/VASP homology domain 1 (EVH1), responsible for mediating Ena localization; a glutamate-rich region; a proline-rich core; and an EVH2 domain, containing G- and F-actin binding sites and a tetramerization motif (Figure 3A). Like Fascin, Ena is required for parallel actin filament bundle formation during S10B (Gates *et al.*, 2009).

**Figure 3.**
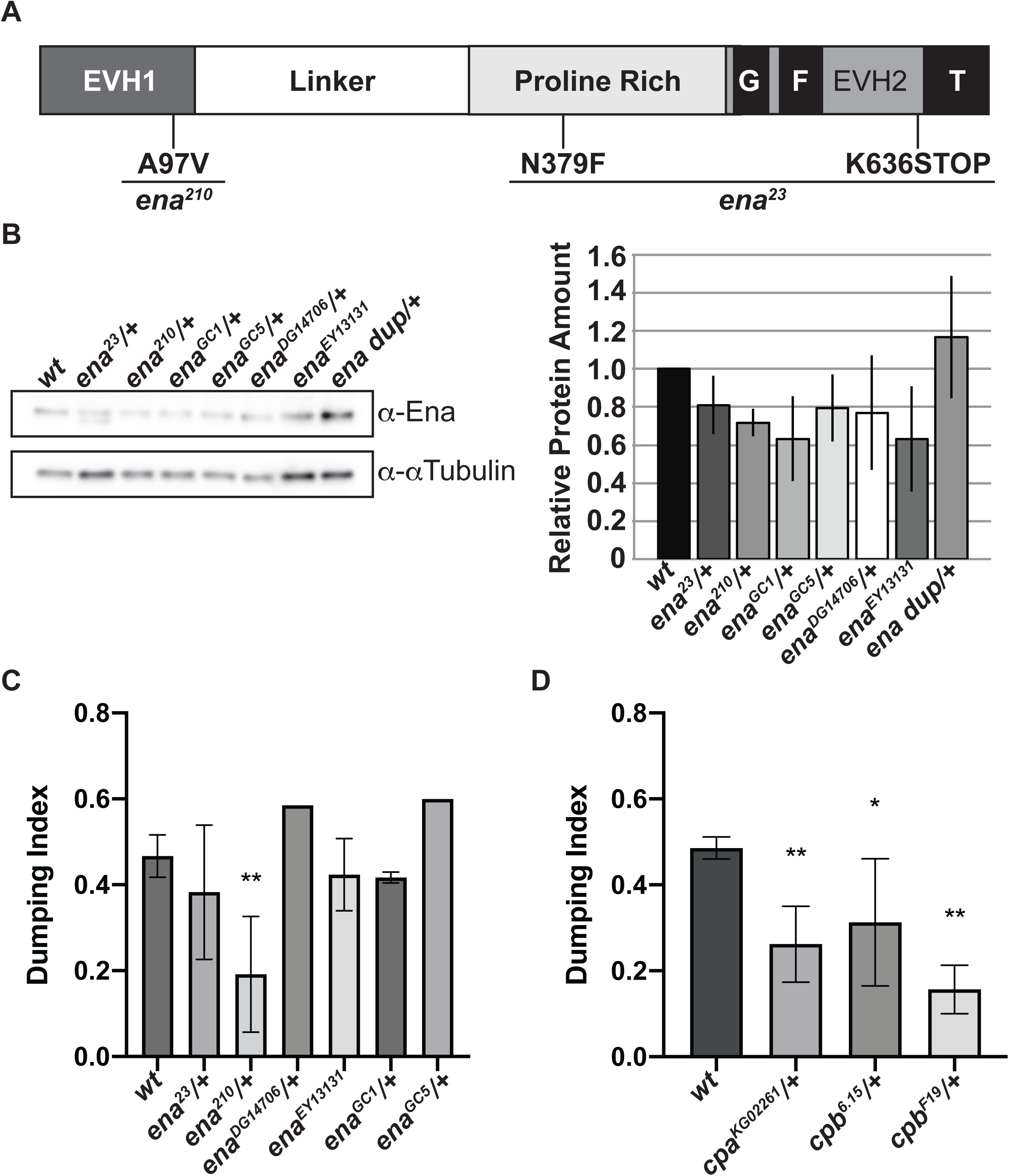
Altering Ena activity strongly sensitized S10B follicles to the effects of COX inhibition. A. Schematic detailing Ena protein structure and the mutations carried by the *ena*^*23*^ and *ena*^*210*^ alleles. B. Representative western blot and quantification of Ena levels for genotypes indicated. C. Graph of pharmaco-genetic interaction data for *ena* alleles. D. Graph of pharmaco-interaction data for *cp* alleles. Ena is a large, multi-domain protein containing an Ena/VASP homology domain 1 (EVH1) and EVH2 domain separated by a poorly conserved glutamine-rich linker region and a conserved proline-rich region; G= G-actin binding domain, F= F-actin binding domain, T= tetramerization domain. *ena*^*23*^ contains a missense mutation (N379F) and one nonsense mutation (K636STOP), resulting in the production of a truncated protein lacking the tetramerization domain (A). *ena*^*210*^ contains a single missense mutation (A97V), which has been shown to disrupt Ena interaction with Zyxin, an Ena interacting partner, *in vitro* (A). Heterozygosity for *ena*^*23*^, *ena*^*210*^, *ena*^*GC1*^, *ena*^*GC5*^, or *ena*^*DG14706*^ or homozygosity for *ena*^*EY13131*^ results in a mild decrease in total Ena levels compared to wild-type controls (B, n=2, error bars = SD; note both full-length and truncated Ena are observed in the *ena*^*23*^/+ lane). Additionally, heterozygosity for an Ena duplication (*ena* dup; Dp(2;2)Cam18/+) results in a mild increase in total Ena levels compared to wild-type controls (B). Heterozygosity for numerous *ena* alleles including 2 null alleles (*ena*^*GC1*^ and *ena*^*GC5*^), a *hobo* element insertion into the *ena* locus (*ena*^*DG14706*^), and a point mutation resulting in ablation of Ena’s EVH2 domain (*ena*^*23*^), as well as homozygosity for a P element insertion into the *ena* locus (*ena*^*EY13131*^), fail to modify follicle sensitivity to the effects of COX inhibition (C). However, heterozygosity for the *ena*^*210*^ allele, which bears a point mutation known to disrupt Ena’s EVH1 domain, significantly enhances the ability of 1.5mM aspirin to inhibit nurse cell dumping compared to wild-type controls (C). Heterozygosity for a P element insertion into the locus of the alpha-subunit of Capping protein (*cpa*^*KG02261*^), and two hypomorphic allele of the beta-subunit of Capping protein (*cpb*^*6.15*^ and *cpb*^*F19*^) significantly enhance the ability of 1.5mM aspirin to block nurse cell dumping compared to wild-type controls (D). n’s are indicated in Table 1. Error bars = SD. ** indicates p <0.0001, * indicates p<0.05.

To assess whether Ena is a downstream target of PG signaling during the actin remodeling occurring during S10B, we asked if reduced Ena levels modified follicle sensitivity to COX inhibition. Here, we took advantage of six different *ena* alleles. These alleles included two well-characterized alleles recovered from a gamma ray-induced mutagenesis screen (*ena*^*GC1*^ and *ena*^*GC5*^), two alleles recovered from insertional mutagenesis screens (*ena*^*DG14706*^ and *ena*^*EY13131*^), and two well-characterized alleles recovered from an ethyl nitroso-urea mutagenesis screen (*ena*^*23*^ and *ena*^*210*^). Both *ena*^*GC1*^ and *ena*^*GC5*^ alleles are null alleles, resulting from inversions within the *ena* locus. Neither *ena*^*DG14706*^ nor *ena*^*EY13131*^ have been extensively characterized, but are predicted to result in reduced Ena levels. The viability of *ena*^*EY13131*^ homozygotes suggests it is likely a weak, hypomorphic allele. The *ena*^*23*^ allele contains one missense mutation (N397F) of unknown functional consequence and a nonsense mutation (K636STOP) resulting in loss of the tetramerization motif within the EVH2 domain (Figure 3A). The *ena*^*210*^ allele contains a missense mutation (A97V) within the EVH1 domain (Figure 3A), which disrupts *in vitro* interaction between Ena and Zyxin (Ahern-Djamali *et al.*, 1998) and likely alters Ena localization.

Using our pharmaco-genetic interaction screen, we find reduced levels of Ena result in an allele-dependent enhancement of follicle sensitivity to the effects of COX-inhibition. Heterozygosity for *ena*^*GC1*^ (0.417, ±0.013 SD, n=2), *ena*^*GC5*^ (0.6, n=1), *ena*^*DG14706*^ (0.585, n=1), and *ena*^*23*^ (0.383, ±0.156 SD, n=5) or homozygosity for *ena*^*EY13131*^ (0.423, ±0.084 SD, n=2) fails to significantly modify follicle sensitivity to COX inhibition compared to wild-type controls (0.467, ±0.049 SD, n=11) (Table 1 and Figure 3C). In contrast, heterozygosity for *ena*^*210*^, a localization defective allele, strongly enhances follicle sensitivity to COX inhibition. We find that heterozygosity for *ena*^*210*^ results in a dumping index of 0.192 (±0.135 SD, n=5) compared to 0.467 (±0.049 SD, n=11) for wild-type controls (p=<0.0001) (Table 1 and Figure 3C).

While multiple alleles fail to show an interaction, the *ena*^*210*^ allele strongly suppresses follicle sensitivity to COX inhibition in our pharmaco-genetic interaction screen. There are a number of potential explanations for this finding. It is possible that this allele-specific interaction could be due to genetic background effects that are specific to the *ena*^*210*^ allele. Given that the *ena*^*23*^ and *ena*^*210*^ alleles were generated in the same genetic background and have both been recombined onto the same FRT bearing chromosome, genetic background effects are unlikely to account for the observed differences. It is more likely the allele-specific interaction we observe is due to the nature of alleles tested. Unlike *ena*^*GC1*^, *ena*^*GC5*^, *ena*^*DG14706*^ and *ena*^*EY13131*^, which are predicted to result in decreased Ena levels, the *ena*^*23*^ and *ena*^*210*^ alleles produce stable mutant protein products. Therefore, while heterozygosity for *ena*^*GC1*^, *ena*^*GC5*^, *ena*^*DG14706*^, or homozygosity for *ena*^*EY13131*^ would be expected to result in decreased Ena levels, heterozygosity for *ena*^*23*^, and *ena*^*210*^ would be expected to result in a mixture of both wild-type and mutant protein. Indeed, western blot analysis reveals that total Ena levels are similarly mildly reduced in *ena*^*GC1*^, *ena*^*GC5*^, *ena*^*DG14706*^, *ena*^*23*^, and *ena*^*210*^ heterozygotes and *ena*^*EY13131*^ homozygotes (∼80% of wild-type controls) (Figure 3B). Because the protein product encoded by *ena*^*23*^ lacks a tetramerization domain, we hypothesize that heterozygosity for *ena*^*GC1*^, *ena*^*GC5*^, *ena*^*DG14706*^, or *ena*^*23*^ have similar net effects—namely, a mild reduction in functional Ena levels. In contrast, we hypothesize heterozygosity for *ena*^*210*^ may result in a stronger effect on Ena function, as the mutant protein would be capable of forming tetramers with wild-type Ena. It is possible that such an interaction may result in a partial dominant-negative effect that more strongly impacts functional Ena levels.

The observation that heterozygosity for the *ena*^*210*^ allele strongly sensitizes follicles to the effects of COX inhibition suggests that PG signaling may regulate parallel actin filament bundle formation during S10B, at least in part, by modulating Ena localization. Supporting this idea, using immunofluorescence and confocal microscopy we previously found Ena localization to the barbed ends of actin filament bundles and nurse cell membranes during S10B is reduced in *pxt* mutants, compared to *wildtype* (Spracklen *et al.*, 2014). This qualitative reduction in localization is not due to altered Ena levels as Ena expression at either the mRNA (Tootle *et al.*, 2011; Spracklen *et al.*, 2014) or protein level (Spracklen *et al.*, 2014) is unchanged in *pxt* mutants. Notably, the extent of the reduction in Ena localization corresponds to the severity of the S10B phenotype in *pxt* mutants, such that when there are nearly no actin filament bundles, there is similarly little to no Ena. As Ena is required for bundle formation/elongation during S10B (Gates *et al.*, 2009), we hypothesize the reduction in Ena localization to sites of bundle formation during S10B is one cause of the bundle defects observed in *pxt* mutants. Together these data suggest PG signaling may be required to promote appropriate Ena localization and/or activity during S10B. Exactly how this regulation is achieved remains unclear, but is likely through post-translational mechanisms. One intriguing means by which PGs may regulate Ena activity is through controlling Fascin. Indeed, Fascin has be shown to promote Ena processivity, increasing filament elongation (Winkelman *et al.*, 2014; Harker *et al.*, 2019). Another mechanism by which PGs may regulate Ena is discussed below.

### Reduced Cp strongly sensitizes follicles to the effects of COX inhibition

The pharmaco-genetic interaction we observed with Ena led us to ask whether reduced levels of Cp, a functional antagonist of Ena/VASP activity (Bear *et al.*, 2002; Bear and Gertler, 2009), also modified follicle sensitivity to COX inhibition. Cp binds to the barbed ends of actin filaments and prevents their elongation by blocking the addition of actin monomers to the growing filament [reviewed in (Cooper and Sept, 2008)]. Cp is a functional heterodimer composed of Capping protein α (Cpa; *Drosophila* Capping protein alpha, Cpa) and Capping protein α (Cpb; *Drosophila* Capping protein beta, Cpb). Additionally, Cp has been shown to play critical roles during *Drosophila* oogenesis, including oocyte determination (Gates *et al.*, 2009), parallel actin filament bundle formation during S10B (Gates *et al.*, 2009; Ogienko *et al.*, 2013), and border cell migration (Gates *et al.*, 2009; Lucas *et al.*, 2013; Ogienko *et al.*, 2013). With these findings in mind, we asked if reduced Cp could alter follicle sensitivity to COX inhibition using a lethal P-element insertional mutation in *cpa* (*cpa*^*KG02261*^) and two EMS hypomorphic *cpb* alleles (*cpb*^*6.15*^ and *cpb*^*F19*^). Initially, we hypothesized that if reduced levels of Ena sensitized follicles to the effects of COX inhibition, then reduction of a negative regulator of Ena activity would have an opposite effect - suppression of follicle sensitivity to COX inhibition. Surprisingly, we found heterozygosity for all alleles of *cpa* and *cpb* tested sensitize follicles to the effects of COX inhibition (Table 1 and Figure 3D). Heterozygosity for *cpa*^*KG02261*^ results in a dumping index of 0.262 (±0.088 SD, n=3) compared to 0.486 (±0.026 SD, n=9) for wild-type controls (p=<0.0001). Heterozygosity for *cpb*^*6.15*^ results in a dumping index of 0.313 (±0.148 SD, n=4; p=0.0043), and heterozygosity for c*pb*^*F19*^ results in a dumping index of 0.157 (±0.056 SD, n=3; p=<0.0001). These data reveal that reduced Cp enhances the effects of COX inhibition to block nurse cell dumping and follicle development.

Based on the *in vitro* antagonistic relationship between Ena and Cp, it seems surprising that reduction of either enhances the effects of COX inhibition. However, loss of Ena and loss of Cp were previously shown to result in similar, but subtly different phenotypes during S10B of oogenesis (Gates *et al.*, 2009). Specifically, loss of either Ena or Cp results in loss of nurse cell cortical actin integrity and leads to impaired nurse cell dumping due to parallel actin filament bundle formation defects (Gates *et al.*, 2009). Whereas loss of Ena results in a substantial decrease in the number of cytoplasmic actin bundles formed during S10B, loss of Cp severely disrupts the organization/distribution of the bundles, but not their overall number (Gates *et al.*, 2009). Based on their data, Gates *et al*. reasoned that capping/anti-capping activity must be carefully balanced to promote appropriate cytoplasmic actin filament bundle formation during S10B (Gates *et al.*, 2009). Our findings suggest that PG signaling may play a role in regulating this balance, thereby promoting appropriate parallel actin filament bundle formation during oogenesis.

### Reduced MRLC strongly sensitizes follicles to the effects of COX inhibition

Non-muscle myosin II mediates the contraction of actin filaments and is comprised of two copies of three subunits: Myosin Heavy Chain (*Drosophila* Zipper, Zip), Myosin Essential Light Chain (*Drosophila* Mlc-c), and MRLC (*Drosophila* Spaghetti squash, Sqh). Phosphorylation of MRLC by Rho kinase (ROCK) and Myosin Light Chain Kinase (MLCK) activates myosin (Tan and Leung, 2009; Vicente-Manzanares *et al.*, 2009). MRLC inactivation is achieved by dephosphorylation via Myosin Phosphatase (MYPT; *Drosophila* Mbs and Flw; (Grassie *et al.*, 2011)).

The pharmaco-genetic interaction screen identified MRLC as a strong enhancer of follicle sensitivity to COX inhibition. We tested three MRLC alleles. *sqh*^*AX3*^ is a null allele derived from a 5kb genomic deletion removing most of the *sqh* locus (Jordan and Karess, 1997). *sqh*^*EY09875*^ is the result of a P-element insertion upstream of the *sqh* translation start site. We also tested a large chromosomal deficiency, Df(1)5D, that removes 51 loci, including *sqh*. As expected all three alleles are homozygous lethal. In our assay, we find heterozygosity for *sqh* enhances follicle sensitivity to COX inhibition, with heterozygosity for *sqh*^*AX3*^ resulting in a dumping index of 0.266 (±0.072 SD, n=3; p=0.0010), heterozygosity for *sqh*^*EY09875*^ resulting in a dumping index of 0.294 (±0.159 SD, n=4; p=0.0118), and heterozygosity for *DF(1)5D* resulting in a dumping index of 0.374 (±0.255 SD, n=3; p=0.2127) (Table 1 and Figure 4A). These data suggest PGs normally promote MRLC activity. MRLC is required for nurse cell contraction during S11, as germline loss of MRLC blocks nurse cell dumping but does not cause actin bundle defects (Wheatley *et al.*, 1995). While loss of Pxt causes severe actin bundle defects, unlike other mutants lacking actin bundles, the nurse cell nuclei do not plug the ring canals; this finding indicates that contraction fails in the absence of Pxt (Tootle and Spradling, 2008; Groen *et al.*, 2012). Given the pharmaco-genetic interaction, we hypothesize PGs activate MRLC to drive nurse cell contraction required for nurse cell dumping.

**Figure 4.**
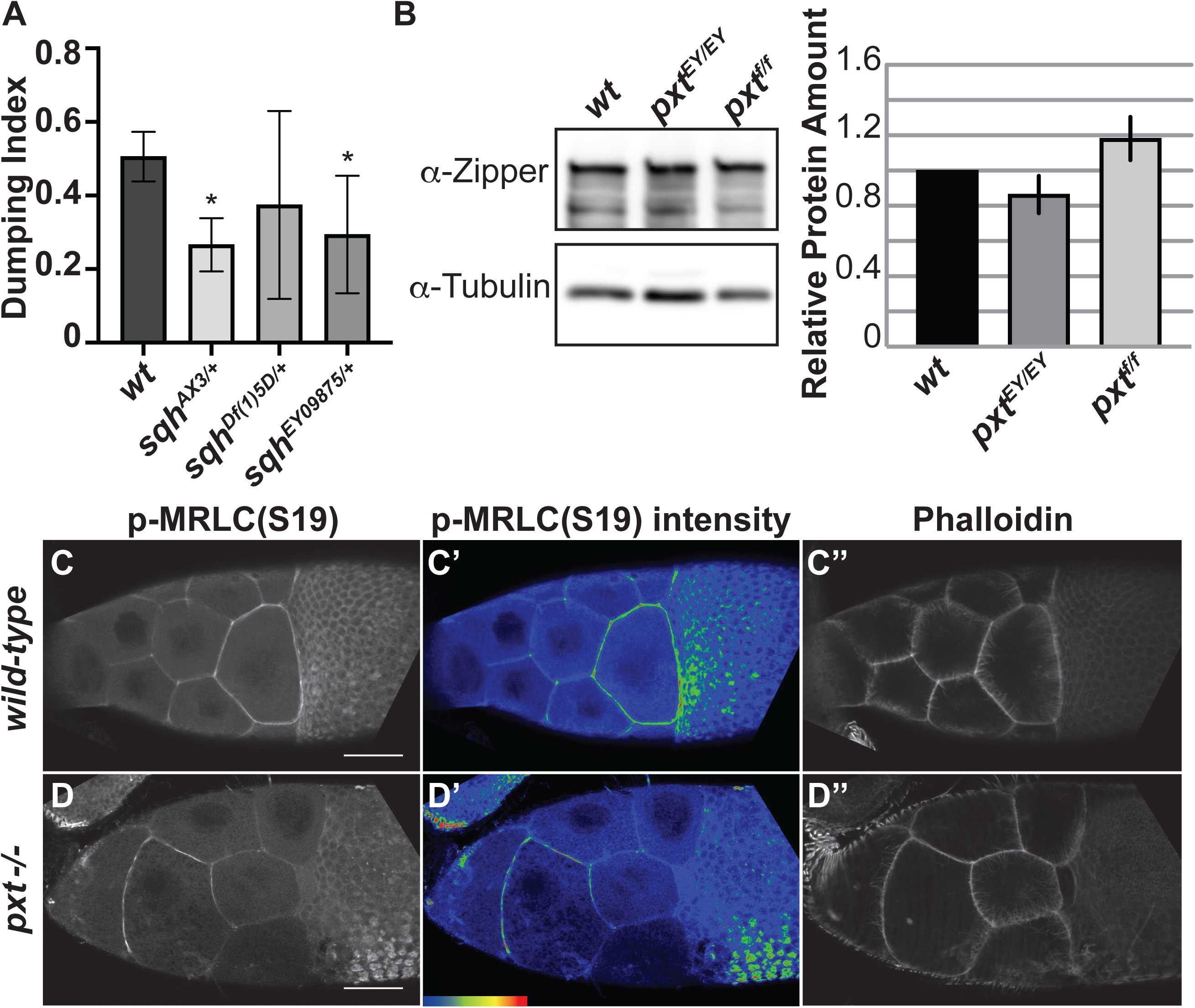
PGs pharmaco-genetically interact with and regulate the activity of MRLC. A. Graph of pharmaco-interaction data for *sqh* alleles; *sqh* encodes the *Drosophila* MRLC. B. Representative western blot and quantification of Zipper (*Drosophila* MHC) levels for genotypes indicated. Tubulin was used as a loading control and Zipper protein levels were normalized to Tubulin. C-D”. Maximum projections of 2-3 confocal slices of S10 follicles of the indicated genotypes. C,D. Phosphorylated myosin regulatory light chain (S19), white. C’,D’. Phosphorylated myosin regulatory light chain (S19) pseudocolored with Rainbow RGB, red indicating the highest intensity pixels. C’’,D’’. Phalloidin (F-actin), white. Heterozygosity for *sqh*^*AX3*^ or *sqh*^*EY09875*^ significantly enhances the ability of aspirin to inhibit nurse cell dumping compared to wild-type controls, and heterozygosity for *sqh*^*Df*(1)5D^ results in a mild increase in aspirin’s ability to inhibit nurse cell dumping (A). Loss of PG signaling using two different Pxt alleles, *pxt*^*EY/EY*^ or *pxt*^*f/f*^, does not alter total Zipper (*Drosophila* MHC) levels (B; n=3, error bars = SD). Loss of PG signaling alters phospho-MRLC localization on nurse cell membranes with the *pxt* mutants have patchy and aberrant enrichment on the anterior nurse cells (C-D’). The change in phospho-MRLC is not due to cortical actin breakdown, as the cortical actin is intact in the *pxt* mutant (C’’, D’’). Error bars = SD. * indicates p<0.05. Scale bars= 50μm.

To explore this idea further, we assessed how loss of Pxt affects the level and activity of non-muscle Myosin II. While immunoblotting for MRLC was unsuccessful, immunoblots for Zipper, the *Drosophila* Myosin Heavy Chain, reveal that non-muscle Myosin II levels are unchanged when PGs are lost (Figure 4B). Next, we examined MRLC by immunofluorescence (Majumder *et al.*, 2012; Aranjuez *et al.*, 2016); MRLC localization is dependent on its activation by phosphorylation (Vicente-Manzanares *et al.*, 2009). During S10B, active MRLC is enriched at the nurse cell membranes, with the strongest staining in the posterior nurse cells (Figure 4C-C’). When Pxt is lost, active MRLC no longer exhibits the posterior enrichment, and it has a patchy appearance at the nurse cell membranes (Figure 4D-D’). This difference is not due to cortical actin defects, as phalloidin-labeled cortical F-actin is present in areas of low active MRLC in *pxt* mutants (Figure 4D” compared to C”). These data suggest PGs regulate non-muscle Myosin II activity to mediate nurse cell contractility.

## Conclusion

The results of this pharmaco-genetic interaction screen support a model in which PG signaling regulates the localization and/or activity of multiple actin binding proteins to coordinate the actin remodeling events during S10B-11 that are required for female fertility. Specifically, our data reveal a number of factors known to both regulate parallel actin filament bundle formation and structure, and to promote nurse cell contractility as new downstream targets of PG signaling during *Drosophila* oogenesis.

Parallel actin filament bundle formation during S10B requires the activity of numerous actin binding proteins, including Profilin (Cooley *et al.*, 1992), Fascin (Cant *et al.*, 1994), Ena (Gates *et al.*, 2009), Cp (Gates *et al.*, 2009; Ogienko *et al.*, 2013), and Villin (Mahajan-Miklos and Cooley, 1994). Reduced levels of Fascin, Ena, and Cp strongly sensitize S10B follicles to the effects of COX inhibition (Table 1 and Figures 2-3). However, not all proteins required for parallel actin filament bundle formation during S10B exhibit an interaction in our pharmaco-genetic interaction screen, as reduced levels of Villin (Groen *et al.*, 2012) or Profilin fail to modify follicle sensitivity to COX inhibition (Table 1 and Figure 2). These data suggest PG signaling regulates a specific subset of actin binding proteins to promote actin filament bundle formation during *Drosophila* oogenesis.

In addition to promoting actin filament bundle formation during S10B, our data also suggest PG signaling is required to promote nurse cell contractility during S11. During nurse cell dumping (S11), the contractile force required to transfer the nurse cell cytoplasm to the oocyte is generated through non-muscle Myosin II-dependent contraction of the nurse cell cortical actin (Wheatley *et al.*, 1995). Reduced levels of MRLC strongly enhance follicle sensitivity to the effects of COX inhibition and loss of Pxt results in decreased active MRLC at the nurse cell membranes (Table 1 and Figures 2, 4). Additionally, ROCK, which is known to promote non-muscle Myosin II activity through phosphorylation of both the MRLC and Myosin Light Chain Phosphatase (Amano *et al.*, 1996), mildly sensitizes follicles to the effects of COX inhibition (Table 1 and Figure 2). Together, these data are consistent with a model in which PG signaling is required during nurse cell dumping to promote nurse cell contractility through positive regulation of non-muscle Myosin II activity.

Future studies are required to elucidate the molecular mechanisms by which PG signaling regulates the localization/activity of the actin regulators uncovered in this screen. These same mechanisms are likely conserved in higher species. Indeed, both increased PG production (Rolland *et al.*, 1980; Chen *et al.*, 2001; Khuri *et al.*, 2001; Gallo *et al.*, 2002; Denkert *et al.*, 2003; Allaj *et al.*, 2013) and actin binding protein expression (Yamaguchi and Condeelis, 2007; Gross, 2013; Izdebska *et al.*, 2018) are associated with increased invasiveness and poor prognosis in multiple human cancers. Given our screen findings, we speculate that PG signaling regulates actin dynamics to drive cancer progression and this is an important area for future investigation.

## Acknowledgements

We thank Stephanie Meyer for her extensive contributions to the screen as an undergraduate researcher, and George Aranjuez and Jocelyn McDonald for the protocol for immunostaining *Drosophila* follicles for active MRLC. We thank the members of the Tootle Lab for helpful discussions and careful review of the manuscript. A.J.S. was supported by the National Institutes of Health (NIH) Predoctoral Training Grant in Pharmacological Sciences T32GM067795, NIH Postdoctoral Training Grant in Integrated Training in Cancer Model Systems T32CA009156, and NIH Ruth L. Kirschstein National Research Service Award Individual Postdoctoral Fellowship F32GM117803. M.C.L. is supported, in part, by the University of Iowa, Graduate College Summer Fellowship, and has previously been supported by the Anatomy and Cell Biology Department Graduate Fellowship. This work was supported by National Science Foundation grant MCB-115827 and NIH grant GM116885 to T.L.T.

